# *Nanoviricide’s* platform technology based NV-CoV-2 polymer increases the half-life of Remdesivir *in vivo*

**DOI:** 10.1101/2021.11.17.468980

**Authors:** Ashok Chakraborty, Anil Diwan, Vinod Arora, Yogesh Thakur, Vijetha Chiniga, Jay Tatake, Preetam Holkar, Neelam Holkar, Bethany Pond

## Abstract

So far, there are seven coronaviruses identified that infect humans and only 4 of them belong to the beta family of coronavirus (HCoV-HKU1, SARS-CoV-2, MERS-CoV and SARS-CoV). SARS family are known to cause severe respiratory disease in humans. In fact, SARS-CoV-2 infection caused a pandemic COVID-19 disease with high morbidity and mortality. Remdesivir (RDV) is the only antiviral drug so far approved for COVID-19 therapy by the FDA. However, the efficacy of RDV *in vivo* is limited due to its low stability in presence of plasma. This is the report of analysis of the non-clinical pharmacology study of NV-CoV-2 (Polymer) and NV-CoV-2-R (Polymer encapsulated Remdesivir) in both infected and uninfected rats with SARS-CoV-2.

Detection and quantification of NV-CoV-2-R in plasma samples was done by MS-HPLC chromatography analyses of precipitated plasma samples from rat subjects.

i. NV-CoV-2-R show RDV peak in MS-HPLC chromatography, whereas only NV-CoV-2 does not show any RDV-Peak, as expected.
ii. NV-CoV-2 polymer encapsulation protects RDV *in vivo* from plasma-mediated catabolism.
iii. Body weight measurements of the normal (uninfected) rats after administration of the test materials (NV-CoV-2, and NV-CoV-2-R) show no toxic effects on them.

Our platform technology based NV-387-encapsulated-RDV (NV-CoV-2-R) drug has a dual effect on coronaviruses. First, NV-CoV-2 itself as an antiviral regimen. Secondly, RDV is protected from plasma-mediated degradation in transit, rendering altogether the safest and an efficient regimen against COVID-19.

## Introduction

The severe acute respiratory syndrome coronavirus 2 (SARS-CoV-2) that causes COVID-19 was declared a global pandemic on March 11^th^, 2020 by the WHO **[1]**. However, the evidence for therapies against this virus is as yet inadequate.

SARS-CoV-2 virus enters host cells by binding to and fusing with the cell membrane receptor, ACE-2, followed by membrane fusion. Once inside, the virus uses the host cell’s machinery to replicate by using the virus’s RNA dependent RNA polymerase (RdRp) for making genome and transcript copies. Among the different strains of the coronavirus, this non-structural protein is unique in structure which makes it a potentially useful drug target. *Sofosbuvir*, a synthetic analogue of nucleosides and nucleotides, inhibits RdRp has become a successful treatment for hepatitis C infection **[2]**.

*Remdesivir (RDV)*, formerly known as GS-5734, is a nucleotide analogue that is claimed to have been originally developed as a treatment against Ebola **[3]**. This drug can also inhibit coronavirus replication by inhibiting RNA polymerases (RdRp4). This compound has shown broad antiviral activity *in vitro* against Middle East respiratory syndrome coronavirus (MERS-CoV), severe acute respiratory syndrome coronavirus 1 (SARS-CoV-1) and SARS-CoV-2 **[4-6]**.

Based on these facts, RDV was approved by the FDA in various clinical trials for the treatment of COVID-19 **[7]**. In animal studies, RDV has been found effective in protecting rhesus monkeys from MERS-CoV infection, when given prior to infection **[8]**. It also protected African green monkeys from Nipah virus, the cause of fatal encephalitis and rhesus monkeys from Ebola virus **[9, 10]**. A randomized, well marked, controlled animal study with 12 rhesus monkeys infected with SARS-CoV-2 reported that an attenuation of respiratory symptoms and reduction in lung damage with RDV administered 12 hours after virus infection **[11]**.

However, efficacy of RDV *in vitro* or in animals does not match with the clinical outcomes in humans. Further, RDV has some side effects. In the Ebola trial, the side effects of RDV were possible liver damage from an increase in liver enzyme levels in the plasma. Similar increases in liver enzymes were found in three U.S. COVID-19 patients were also documented after RDV treatment. Other typical antiviral drug side effects include Nausea and Vomiting **[12];** and also affects kidney and mitochondria **[13, 14]**.

The efficacy of RDV *in vivo* is limited due to the low stability in the plasma. We have tested the stability of RDV encapsulated with our platform technology based polymer NV-CoV-2-R, in presence of plasma *in vitro*, and the result is very supportive that polymer protects RDV from plasma-mediated catabolism **[15]**.

In this paper, we wanted to extend our *in vitro* experiments in an *in vivo* rat model of systemic exposure of NV-CoV-2 and NV-CoV-2-R once per day for 5 days (0, 1, 3, 5, and 7) over a 7-day time period. We compared our results with commercially available Gilead RDV and used DMSO as a negative control. Here we reported our results.

## Materials and Methods

### Test Articles

I. NV-CoV-2 used in Batch ID NV1067-387: Polymer
II. NV-CoV-2-R used in Batch ID NV1067-387-R: Polymer encapsulated RDV
III. NV1067-376: RDV in SBECD (Gilead)

### *In vivo* Treatment

Thirty-six Sprague Dawley rats (Taconic Biosciences, USA) (three/ each sex in control and in treatment groups) were administered with NV-CoV-2 or NV-CoV-2-R, once per day for 5 days (0, 1, 3, 5, and 7) over a 7-day time period. DMSO was used as a vehicle for the control group. Injection on day 0 is considered as a 1^st^ injection and on day 7 is the 5^th^ injection. NV-376 (RDV-in-SBECD; Gilead) was given as two doses on day 1 followed by a daily doses through day 7. Each compound was delivered via slow-push IV injection. Blood samples for systemic exposure assay were collected at 0, 0.08, 0.5, 1, 2, 4, 8 and 24 hours after 1^st^ and 5^th^ injection of the drugs.

Blood samples were taken from one animal per sex in each test article treated group at each time point. Same animal was used to draw blood across all the time points after “day 0” and “day 7” injections.

The procedures for *in vivo* experiments were done by Dr. Krishna Menon from AR Biosystems, (17633 Gunn Highway, Odessa, FL 33556), based on the protocol #IACUC No. 14/17ARB. The study design are shown below in **Table-1**.

**Table-1:**
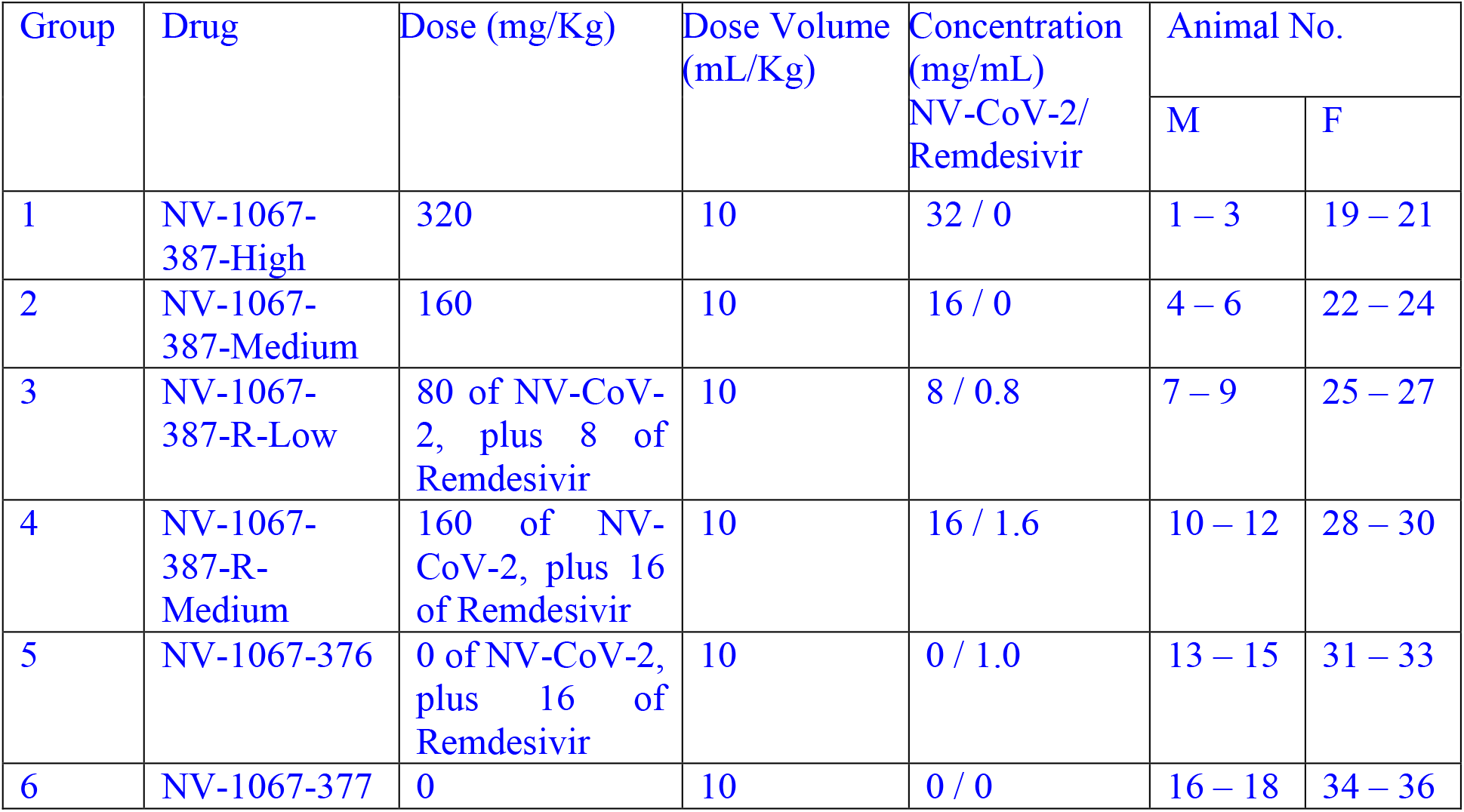
Study Design.

| Group | Drug | Dose (mg/Kg) | Dose Volume (mL/Kg) | Concentration (mg/mL)<br>NV-CoV-2/<br>Remdesivir | Animal No. |  |
| --- | --- | --- | --- | --- | --- | --- |
|  |  |  |  |  | M | F |
| 1 | NV-1067-387-High | 320 | 10 | 32 / 0 | 1 – 3 | 19 – 21 |
| 2 | NV-1067-387-Medium | 160 | 10 | 16 / 0 | 4 – 6 | 22 – 24 |
| 3 | NV-1067-387-R-Low | 80 of NV-CoV-2, plus 8 of Remdesivir | 10 | 8 / 0.8 | 7 – 9 | 25 – 27 |
| 4 | NV-1067-387-R-Medium | 160 of NV-CoV-2, plus 16 of Remdesivir | 10 | 16 / 1.6 | 10 – 12 | 28 – 30 |
| 5 | NV-1067-376 | 0 of NV-CoV-2, plus 16 of Remdesivir | 10 | 0 / 1.0 | 13 – 15 | 31 – 33 |
| 6 | NV-1067-377 | 0 | 10 | 0 / 0 | 16 – 18 | 34 – 36 |

### Assay for RDV in Plasma by LC-MS Spectroscopy

i. Preparation of standard curve **of** pure RDV (Purchased from Sigma-Aldrich Co. USA). Reagents and their sources are shown in **Table-2**.

**Table-2:**
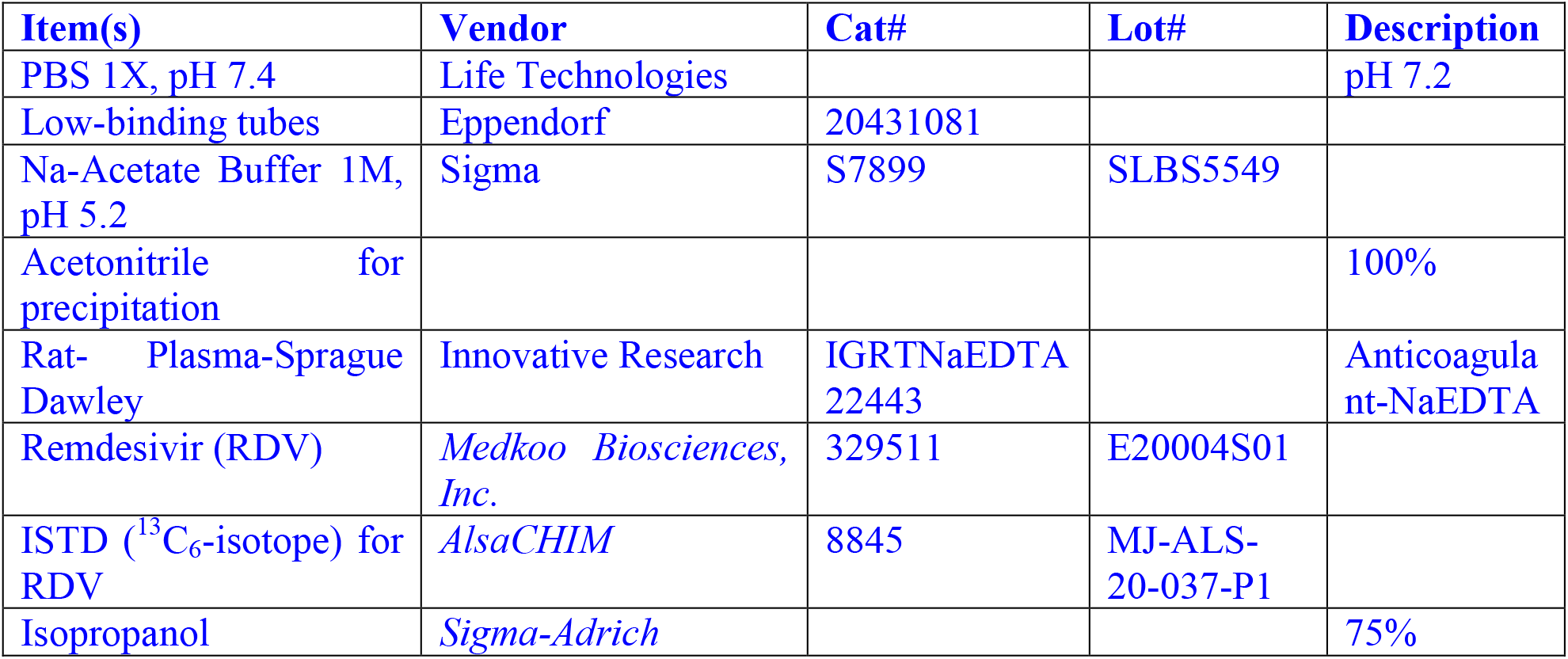
Sources of Reagents: Standard curve preparation of RDV was determined by using LC-MS. Different concentrations of the standard solutions in DMSO + MeOH (1:9) ranging from 0-5 ng/uL final concentrations was prepared. Plasma and/or PBS were used in the reaction mixture. As an internal standard (ISTD), ^13^C_6_ –RDV was used in the mixture. The final concentration of the ISTDs in the reaction mixture becomes 0.125 ng/uL. Extraction of RDV for LC-MS assay was done by using an Acetonitrile cocktail solution (Acetonitrile: ISTDs: 75% Isopropanol at 10:1:1), added at the ratio of 1:4 by volume. The mixture was mixed well and centrifuged for about 5 minutes at 10,000 RPM to separate the precipitated solids from the supernatants. Concentrations of RDV compared to the ratio with ISTDs (^13^C_6_-RDV) was used to generate a standard curve for RDV, as shown in the **result section**.
ii. **Assay of Remdesivir in Rat Plasma after the drugs injection:** To test the *in vivo* stability of our in-house made anti-COVID-19 product, 387 polymer-encapsulated RDV, we have injected our sample along with the polymer only as a negative control, at two different doses. As a positive control, we included other market available Gilead RDV compounds in our in vivo experiments with male and female rat. After the 1^st^ and 5^th^ injection (day 1 and day 7) of the test materials, blood samples were collected at different time points for 24 hrs. Received samples from the test site were pre-diluted (1:2) and diluted further to (1:20) with plasma : 3M Na-acetate, pH 5.0 (1:1). In those samples, reaction mix including ISTDs, 75% Isopropanol and Acetonitrile (1: 1:10) was added at the ratio of 1:4 by volume for the extraction of RDV from the test materials. Vortex and spin the solution for about 5 minutes at 10,000 RPM to separate the precipitated solids from the supernatants.
iii. **Detection of RDV by LC-MS Spectroscopy (as detailed below):** Analysis was performed using the LC and MS analysis conditions shown in **Table-3**, and the multiple reaction monitoring (MRM) data acquisition parameters shown in **Table-4**. Shim-Pack Sceptor™ C18-120 (50 mm x 2.1 mm I.D., 1.9uM) was used as the analytical column.

**Table-3:** Chromatographic Conditions:

| **LC Analysis Conditions** |  |
| --- | --- |
| **HPLC Method:** | RDVmetIS_LC_NoSlit-0.4mL-ExtRinse_202010231219.1cm |
| **Column:** | Shimadzu, Shim Pack, Sceptor, C18-120, 2.1x50 mm, 1.9 um, Cat#227-31120-01 |
| **Pre-Column:** | Guard Column, Shimadzu, Shim-pack, C18-120 EXP (G), 1.9 um, 2.1 x 5 mm, or Restek prefilter: Ultrafield UHPLC, 0.2 um frit, Catalog# 25810. |
| **Mobile Phase:** | A: Water containing 0.05% formic acid<br>B: Acetonitrile containing 0.05% formic acid |
| **Gradient:** | 0-0.3 min: 5% B,<br>0.3-0.35 min: 30% B,<br>0.35-1.5 min: 70% B,<br>1.5-1.8 min: 90% B,<br>1.8-2.8 min: 90% B,<br>2.8-2.9 min: 5% B,<br>2.9-4.5 min: 5% B,<br>(A: Solvent A; B: Solvent B) |
| **Injection Volume:** | 1 uL |
| Auto-sampler was used in external rinsing mode, with R0 = 95/5 Water / Acetonitrile and R3 = 1 / 1 IPA / Methanol. |  |

**Table-4:**
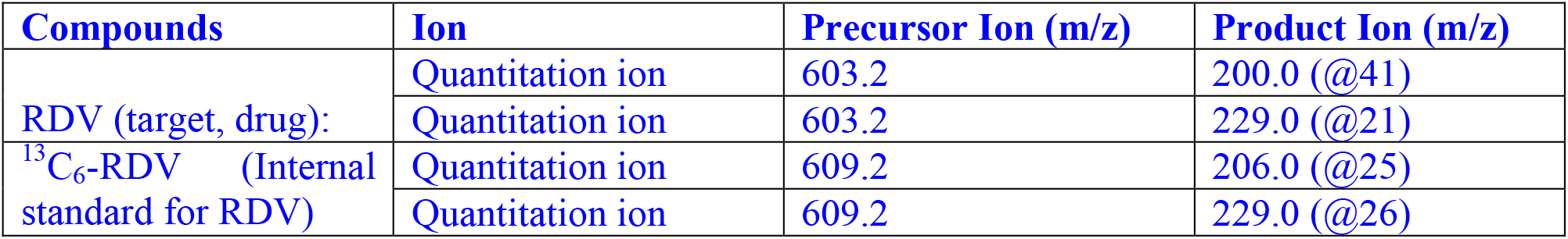
MS MRM Transitions observed (for quantitation):
iv. **Calculation:** From the chromatogram the ratio of RDV and its isotope, ^13^C_6_-ISTD (as an internal standard) was calculated. The amount of RDV was determined using the linear equation derived from their respective standard curve.
v. **Normalization of the value using the dilution factor from the original plasma sample (Table-5):**

| **Compounds** | **Ion** | **Precursor Ion (m/z)** | **Product Ion (m/z)** |
| --- | --- | --- | --- |
| RDV (target, drug): | Quantitation ion | 603.2 | 200.0 (@41) |
|  | Quantitation ion | 603.2 | 229.0 (@21) |
| <sup>13</sup>C<sub>6</sub>-RDV (Internal standard for RDV) | Quantitation ion | 609.2 | 206.0 (@25) |
|  | Quantitation ion | 609.2 | 229.0 (@26) |

**Table-5:** Sample Dilution Calculation.

| **Table-5: Sample Dilution Calculation** |  |  |  |  |
| --- | --- | --- | --- | --- |
| **Samples** | **Received as** | **Further Diluted with the Vehicle** | **Diluted in the reaction Mix** | **Final Dilution Factor** |
| Plasma Samples | 1 : 2 (x 2) pre-diluted | 1 : 20 (x 20) | 1 : 4 (x 4) | 2 x 20 x 4 (x160) |

### Results and Analysis of the Data

(i) A standard curve for RDV from a representative experiment were shown in Table-6 and Fig. 1.

**Fig. 1:**
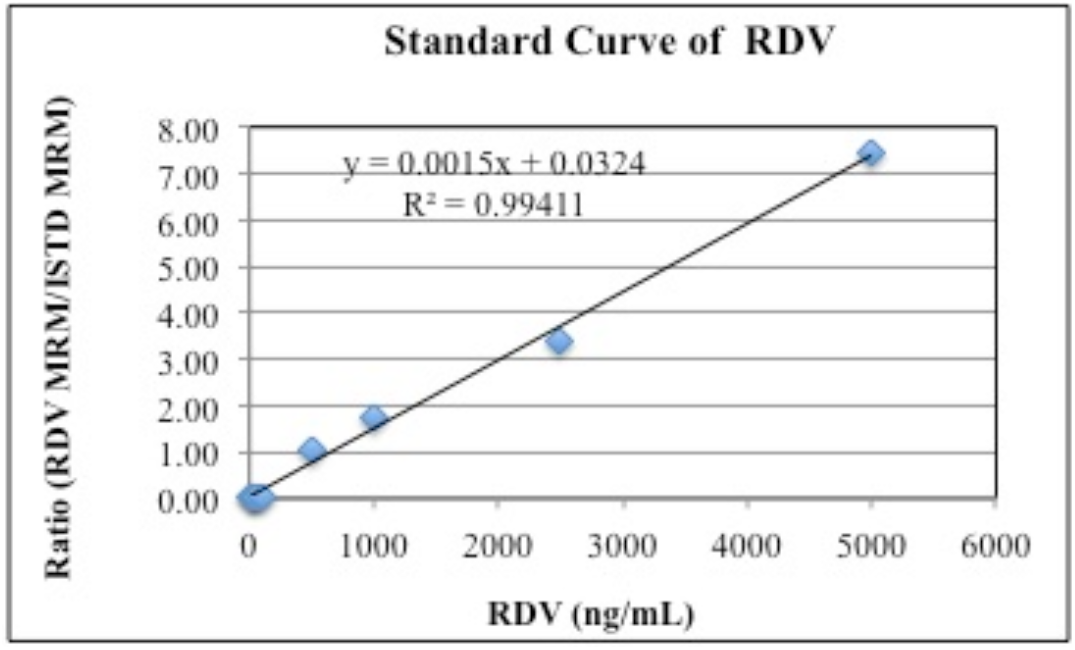
Standard curve of RDV was determined by using LC-MS, using different concentration of the standards solution in DMSO + MeOH (1:9). Final concentrations of RDV ranged from 0-5 ug/mL, which is described in the method section. Values (Mean ± SD) are from a representative experiment done in duplicate.

**Table-6:**
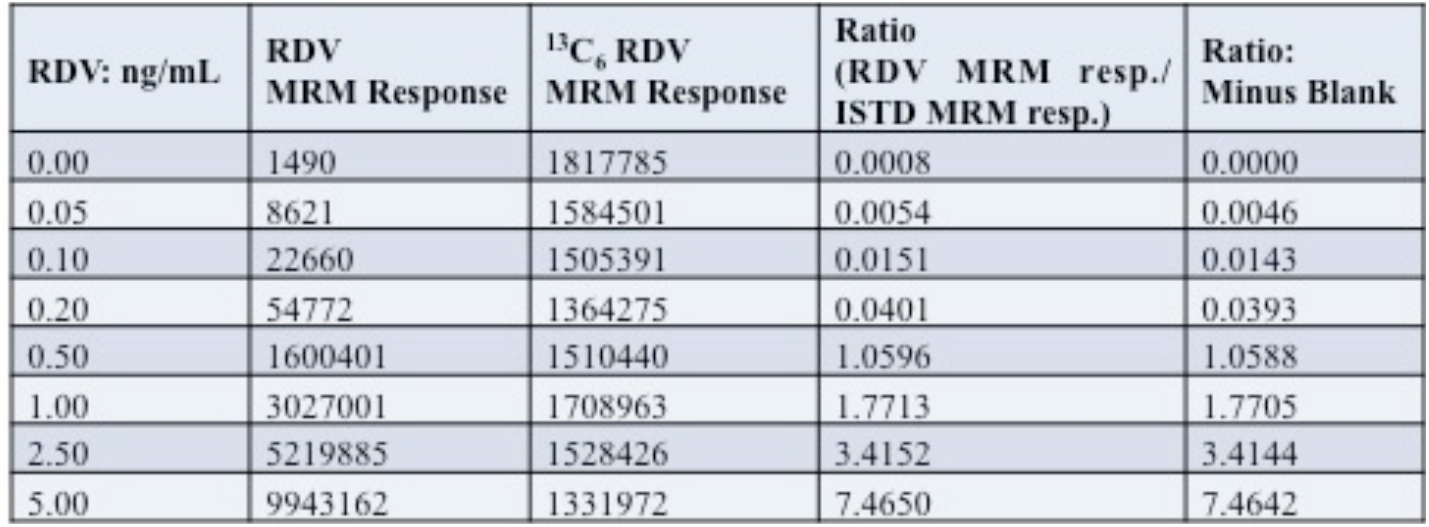
Standard Curve of RDV: A Standard Curve for RDV from a representative experiment
(ii) RDV values in male rats plasma obtained (mg/mL) after 1^st.^ and 5^th^ injection of the drugs were normalized by dividing with the amount of RDV administered (mg/kg of rat body weight) and shown in Fig. 2 and Fig. 3.

| RDV: ng/mL | RDV MRM Response | <sup>13</sup>C<sub>6</sub> RDV MRM Response | Ratio (RDV MRM resp./ ISTD MRM resp.) | Ratio: Minus Blank |
| --- | --- | --- | --- | --- |
| 0.00 | 1490 | 1817785 | 0.0008 | 0.0000 |
| 0.05 | 8621 | 1584501 | 0.0054 | 0.0046 |
| 0.10 | 22660 | 1505391 | 0.0151 | 0.0143 |
| 0.20 | 54772 | 1364275 | 0.0401 | 0.0393 |
| 0.50 | 1600401 | 1510440 | 1.0596 | 1.0588 |
| 1.00 | 3027001 | 1708963 | 1.7713 | 1.7705 |
| 2.50 | 5219885 | 1528426 | 3.4152 | 3.4144 |
| 5.00 | 9943162 | 1331972 | 7.4650 | 7.4642 |

**Fig. 2:**
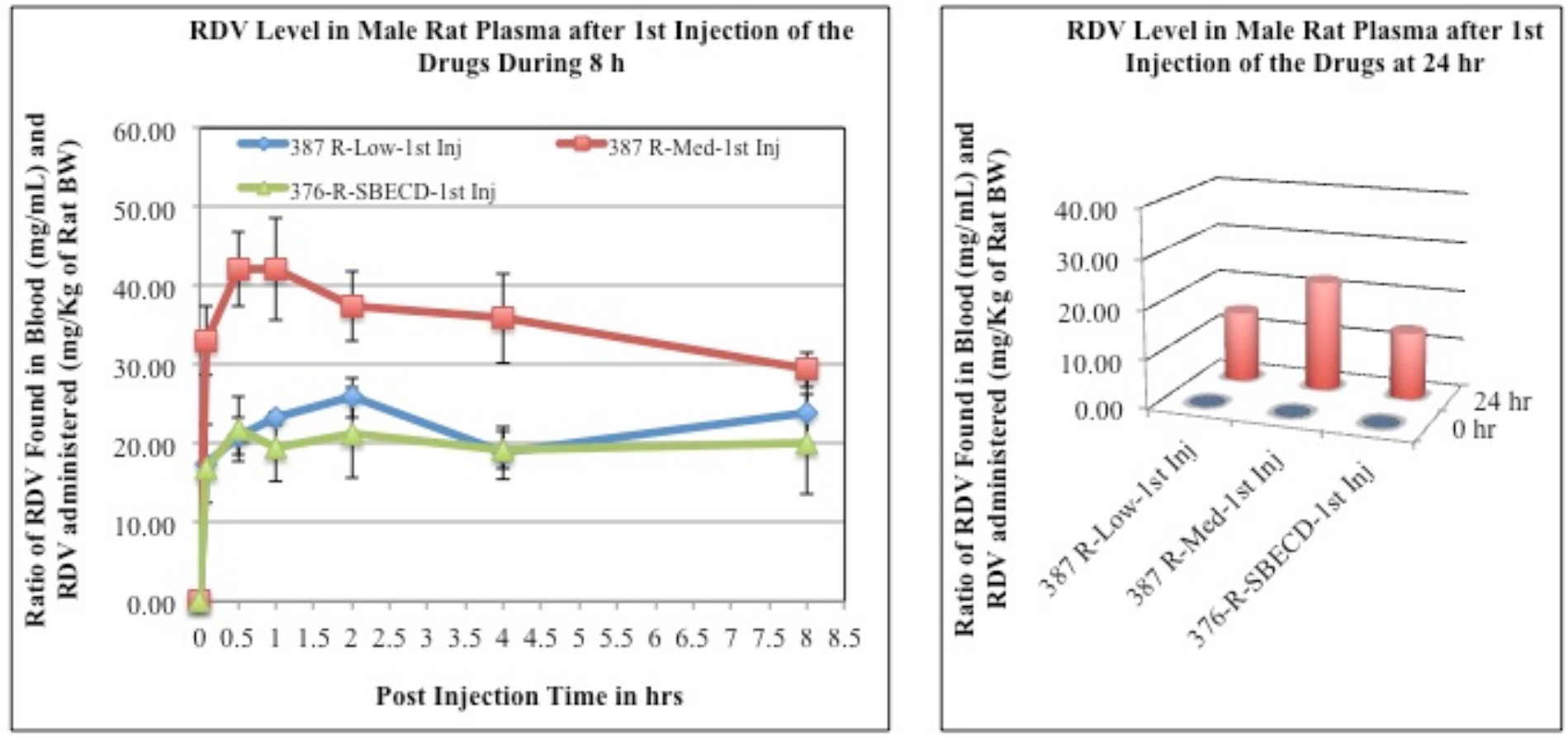
RDV values in **male rat** plasma after **1**^**st**^ **injection** of the drugs. The blood samples collected at different time points after drug adninistration *i*.*v*. to the animals, were collected and measured for RDV level as described in the method section. The values obtained as mg/mL were normalized by dividing with the RDV amount administered (mg/kg of rat body weight). Each data point is the mean (±SD) of 3 values and the experiment was repeated three times with similar results.

**Fig. 3:**
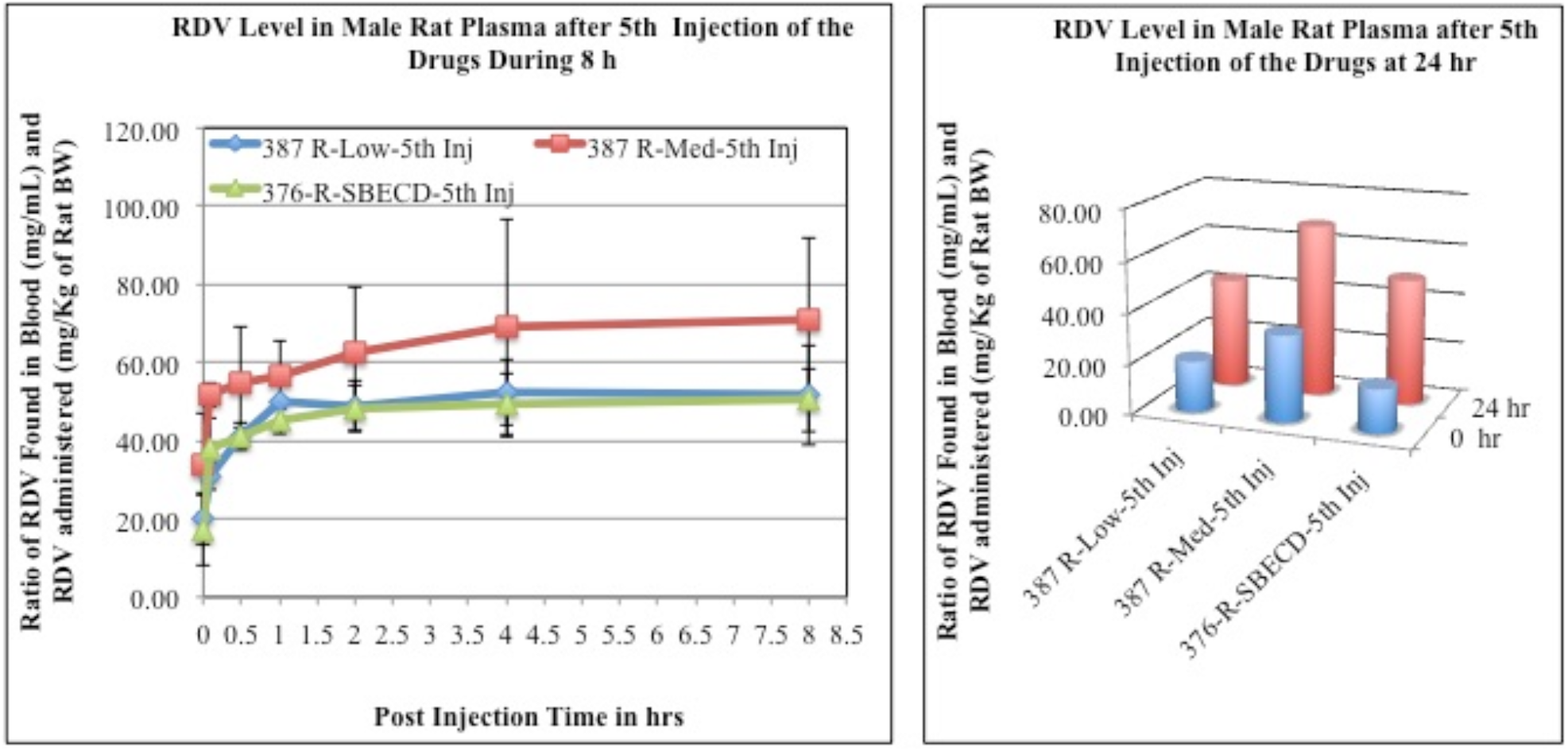
RDV values in **male rat** plasma after **5**^**th**^ **injection** of the drugs. As in Fig. 2.
(iv) RDV values in female rats plasma obtained (mg/mL) after 1^st.^ and 5^th^ injection of the drugs were normalized by dividing with the amount administered (mg/kg of rat body weight) and shown in Fig. 4 and Fig. 5.

**Fig. 4:**
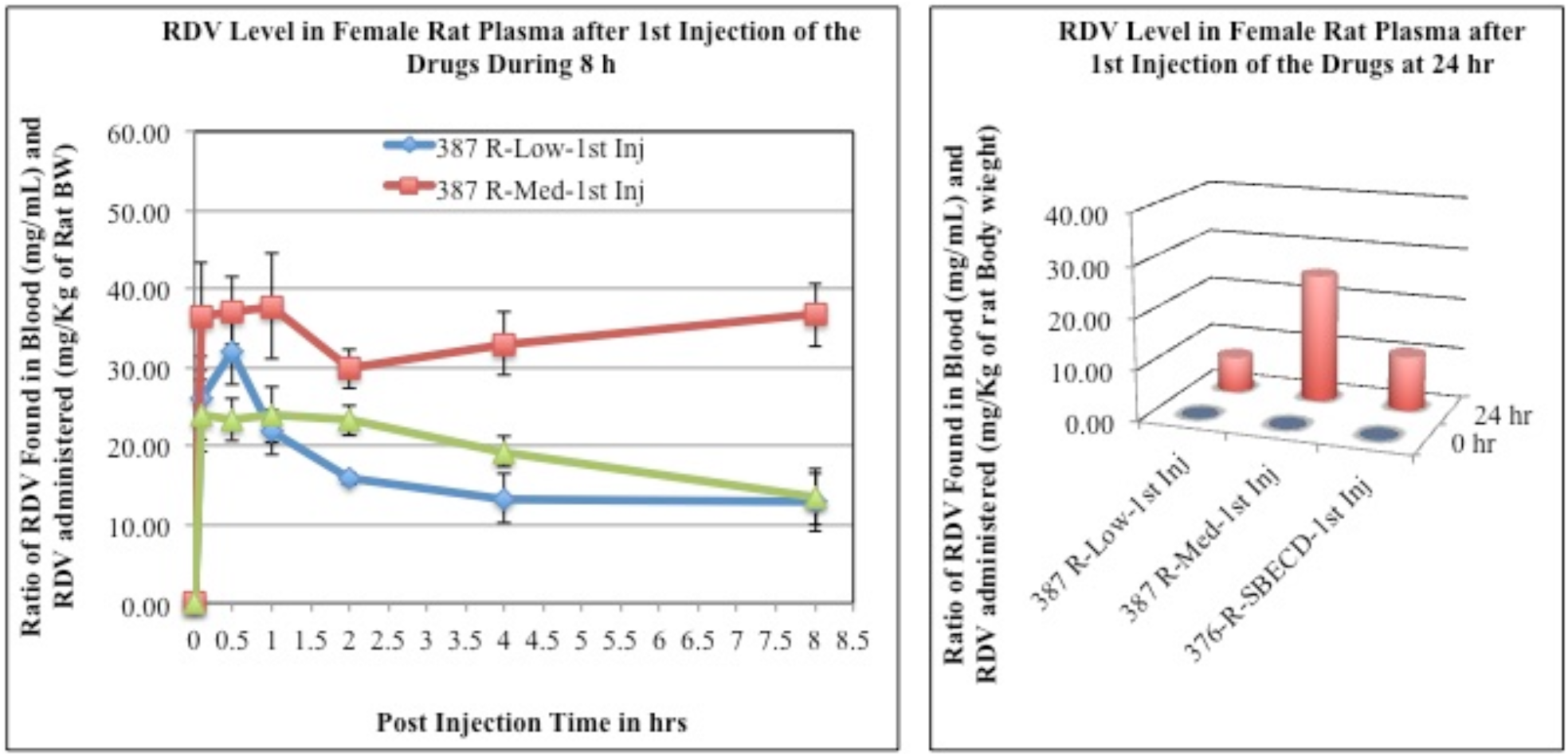
RDV values in **female rat** plasma after **1**^**st**^ **injection** of the drugs. As in Fig. 2.

**Fig. 5:**
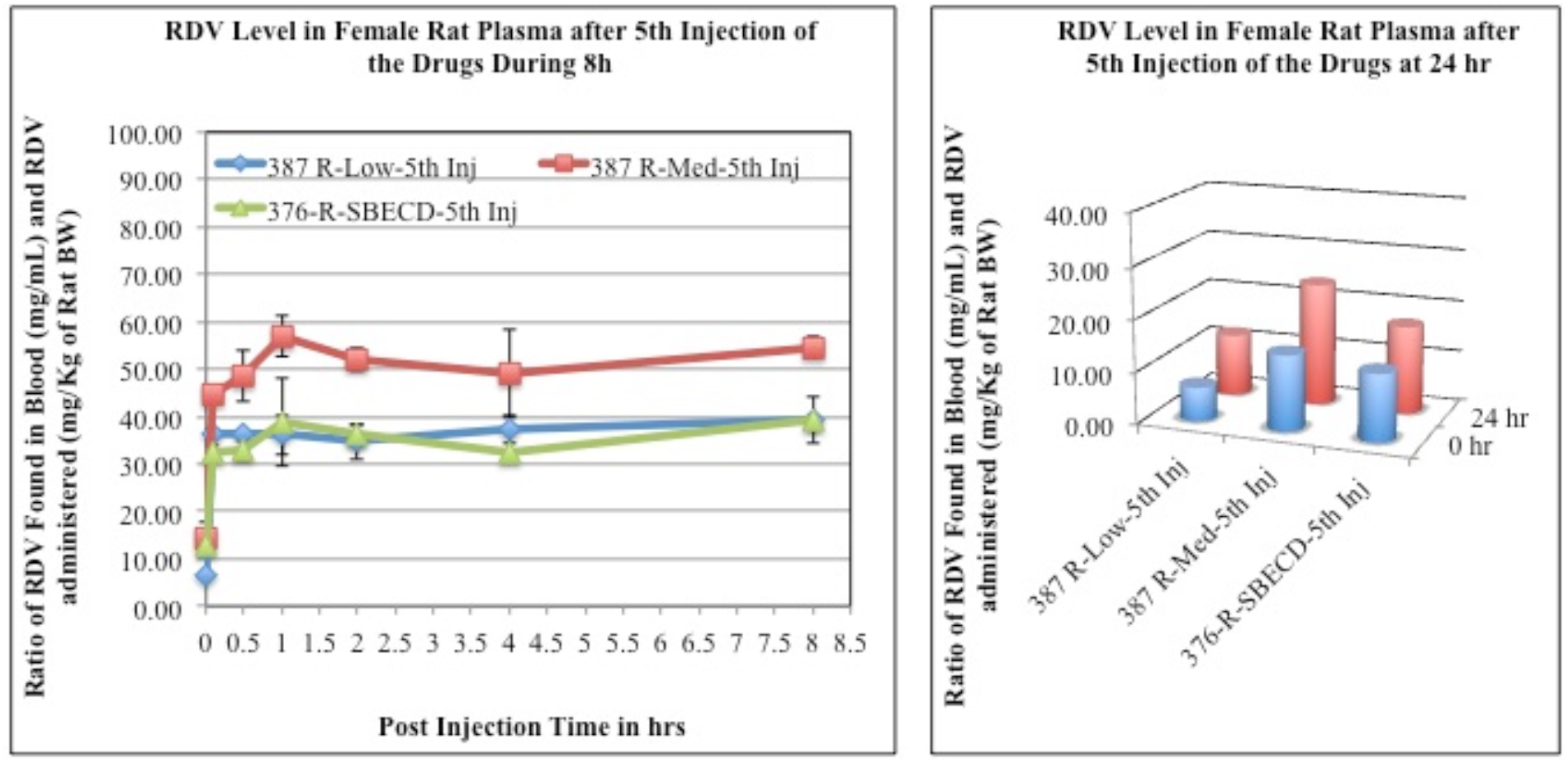
RDV values in **female rat** plasma after **5**^**th**^ **injection** of the drugs. As in Fig. 2.
(v) Comparative analysis of RDV level in male rats plasma after 1^st^ and 5^th^ injection of the Drugs were shown in (Fig. 6). Values were normalized as a ratio of RDV found in blood (mg/mL) and amount of RDV was administered (mg/Kg of rat body weight).

**Fig. 6:**
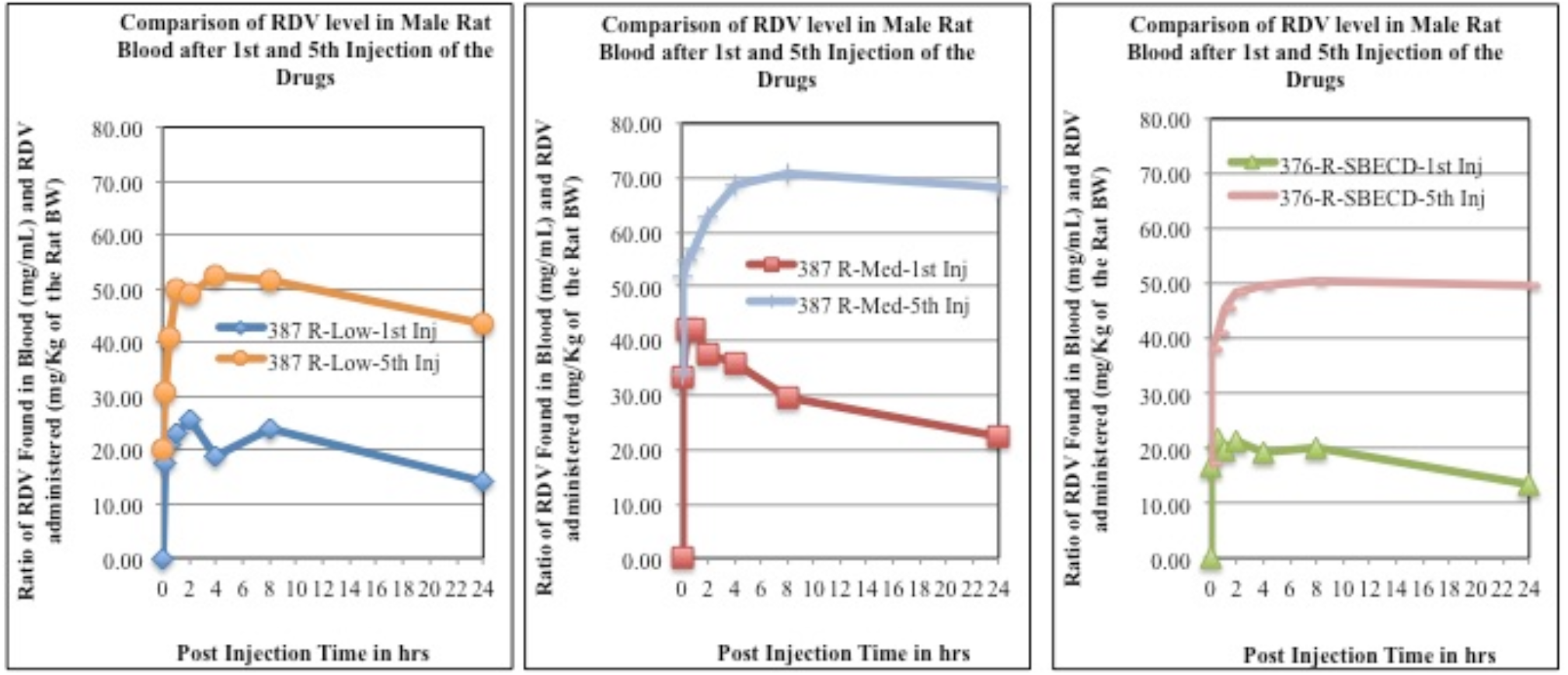
Comparison of RDV level in male rat plasma after 1^st^ and 5^th^ injection of the drugs. As in Fig. 2.
(vi) Comparative analysis of RDV level in female rats plasma after 1^st^ and 5^th^ injection of the Drugs were shown in **Fig. 7**. Values were normalized as a ratio of RDV found in blood (mg/mL) and amount of RDV was administered (mg/Kg of rat body weight).

**Fig. 7:**
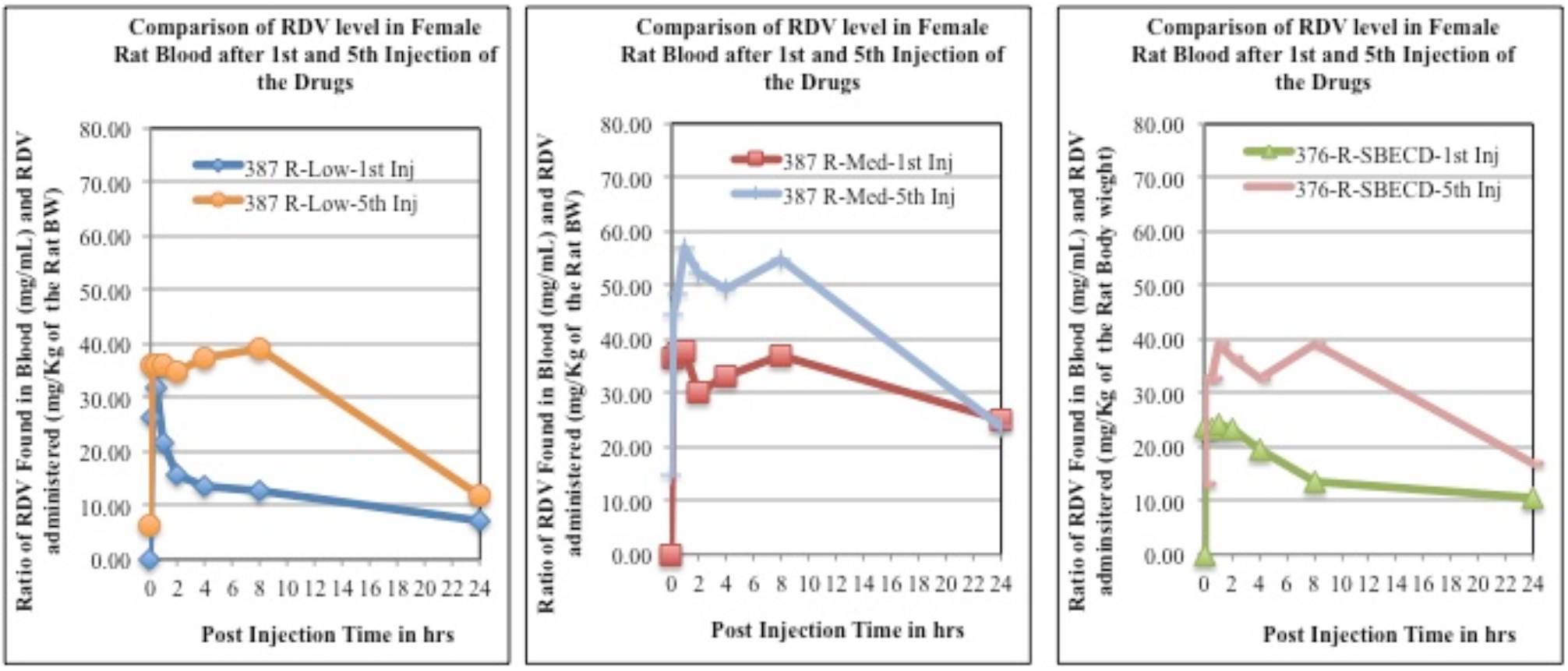
Comparison of RDV level in female rat plasma after 1^st^ and 5^th^ injection of the drugs. As in Fig. 2.
(vi) Comparative analysis of RDV level in male and female rats plasma after 1^st^ injection of the drugs were shown in Fig. 8.

**Fig. 8:**
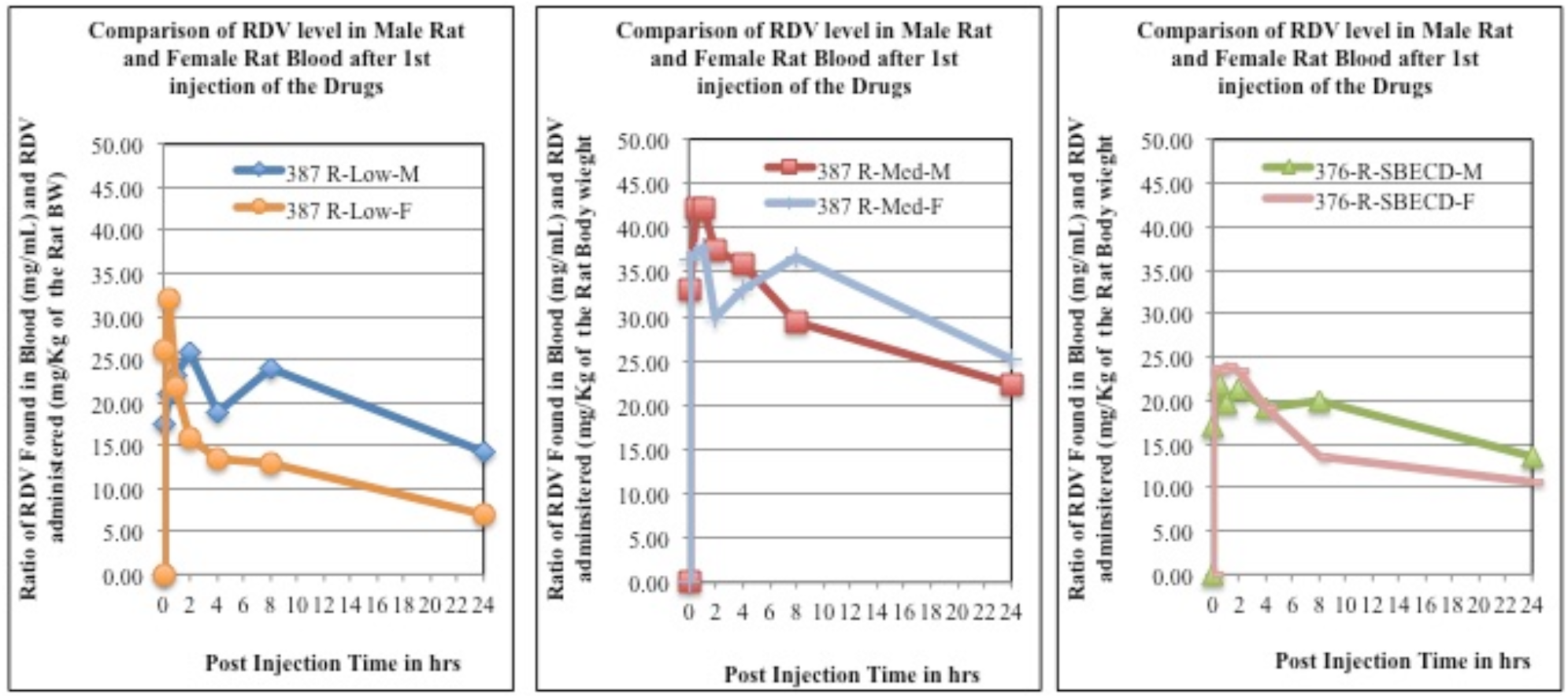
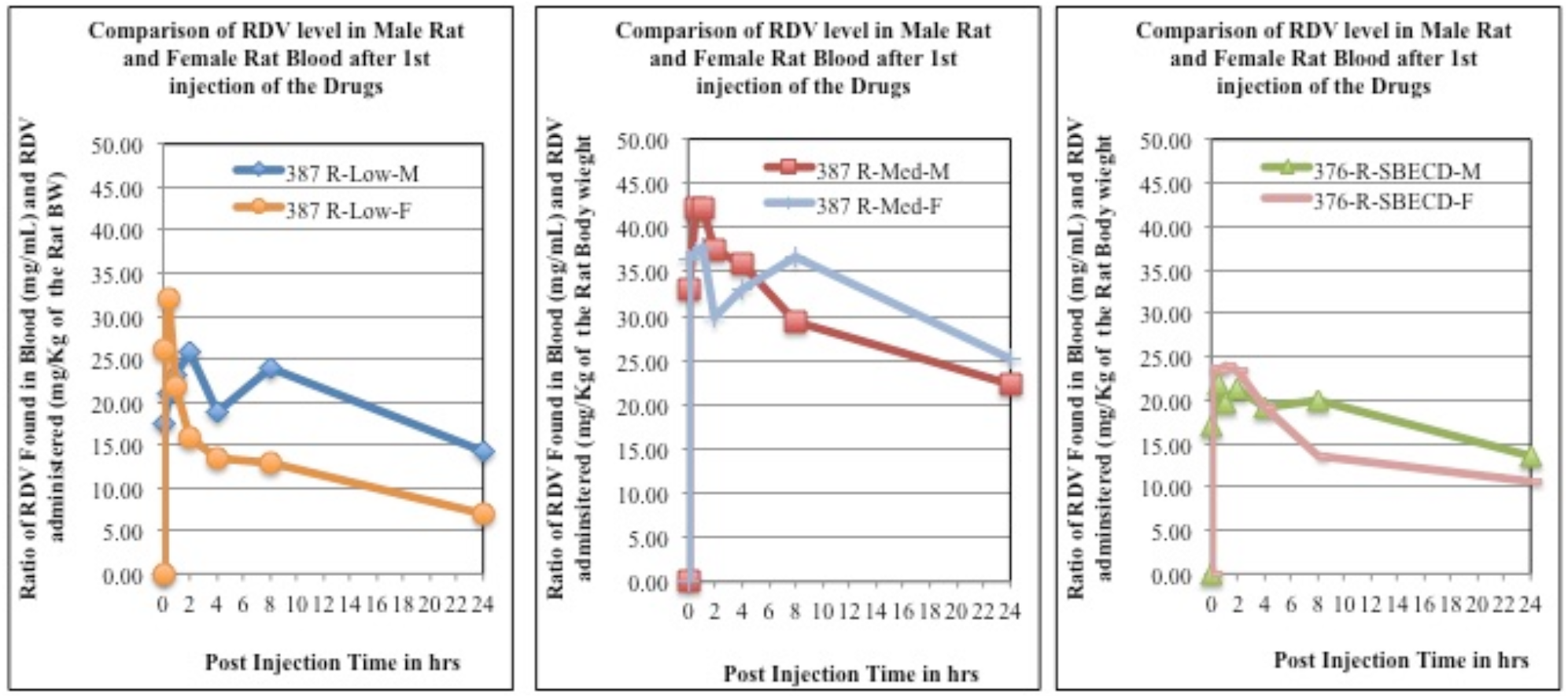
Comparison of RDV level in male and female rat plasma after 1^st^ injection of the drugs
(vii) Comparison of RDV level in male and female rats plasma after 5^th^ injection of the drugs were shown in Fig. 9.

**Fig. 9:**
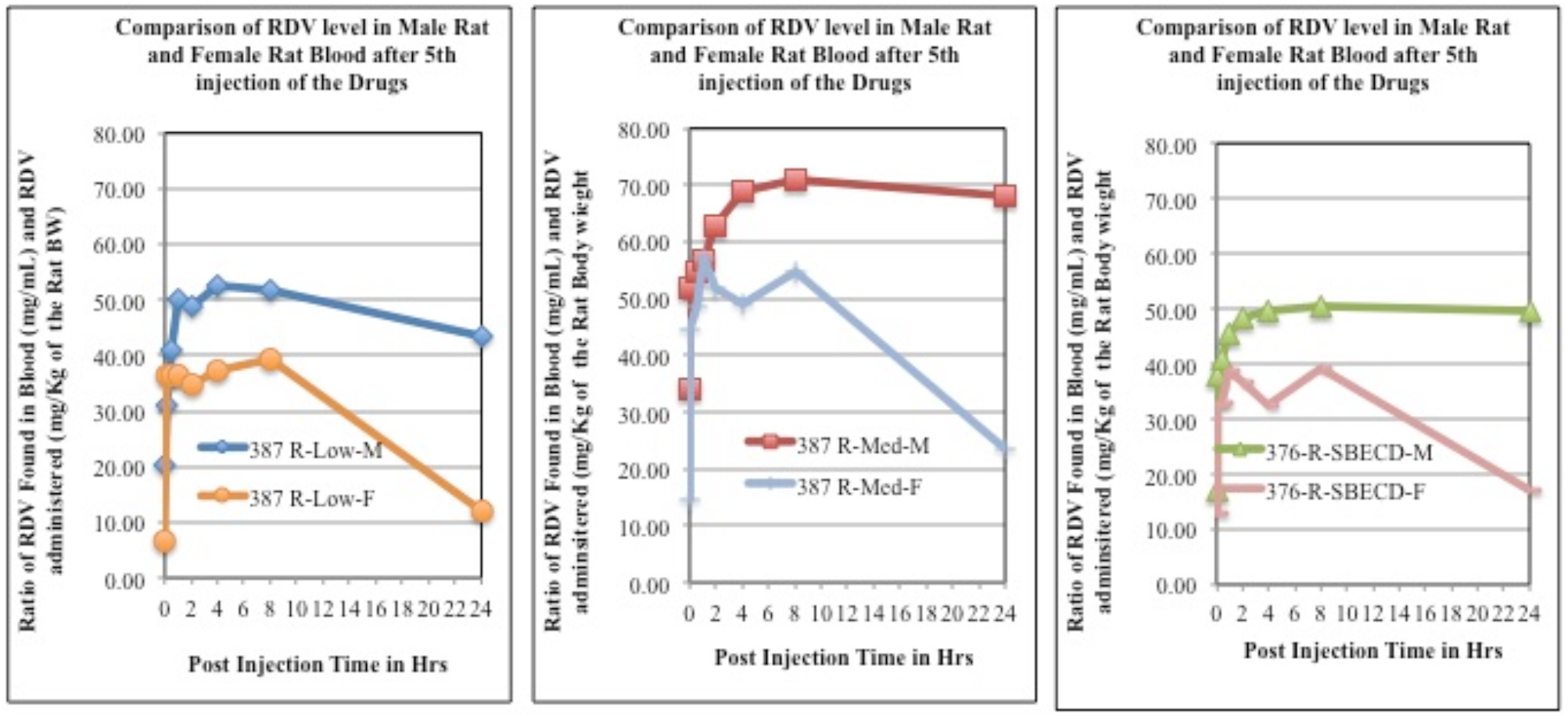
Comparison of RDV level in male and female rat plasma after 5^th^ injection of the drugs
(viii) Toxicity/Tolerability study of NV-CoV-2-R: Loss of body weight analysis of male and female rats after drug administration *i*.*v*. (Fig. 10):

**Fig. 10:**
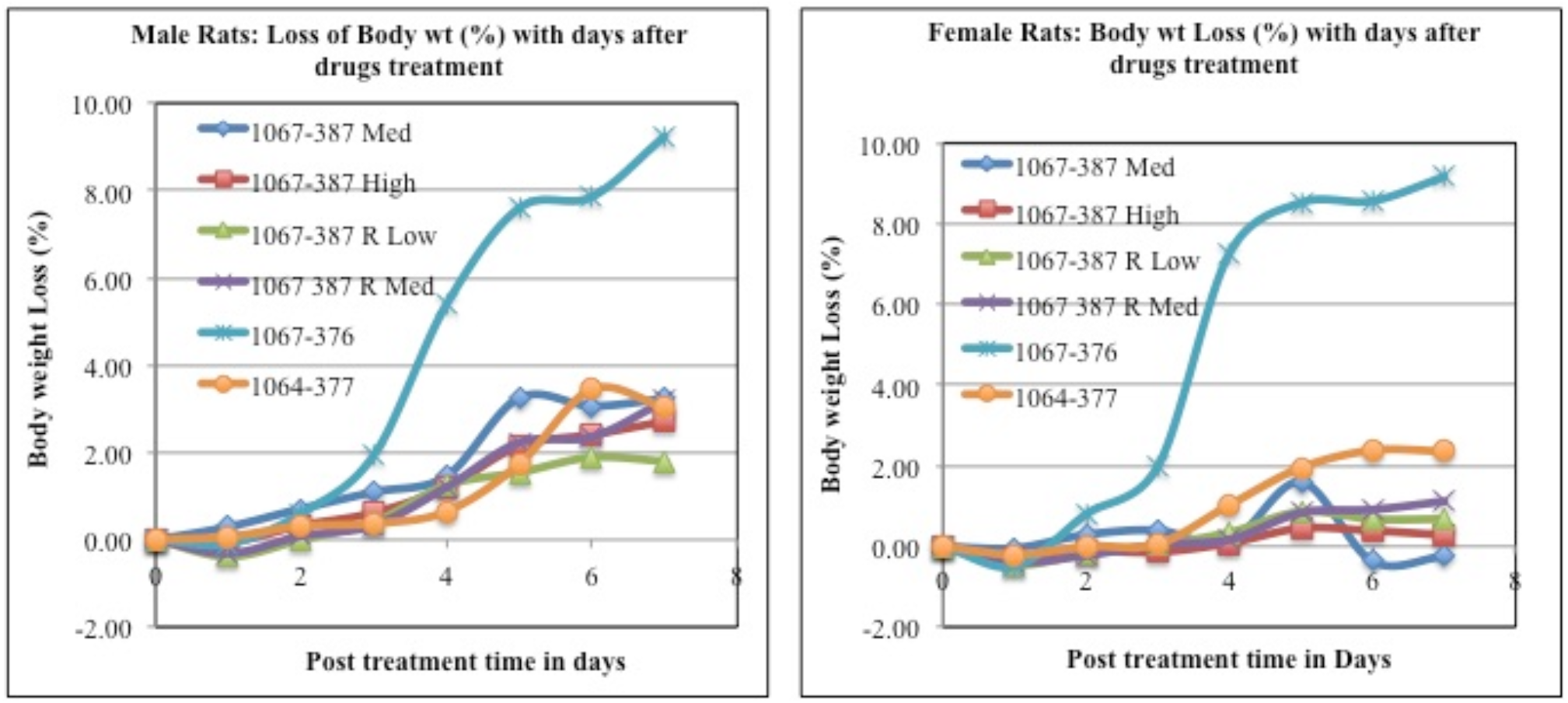
Toxicity/Tolerability study of NV-CoV-2-R: Loss of Body weight analysis of male and memale rats after drug administration i.v.

From all the above figures it appears that:

1. After the 1^st^ injection of 387-R-Low, the accumulation of RDV is better in female rats than in male rats. However, after 5^th^ injection, the differences between male and female rats are insignificant.
2. After the 1^st^ injection of 387-R-Med, the accumulation of RDV is little greater in female rats compared to male rats, which is more evident after 5^th^ injections.
3. No such differences were found in the accumulation of RDV from 376-R-SBECD injections, 1^st^ and 5^th^, both, in male and female rats.
4. There are no differences in RDV accumulation when injected 387-R-Low in male rats, after 1^st^ or 5^th^ injections, either.
5. RDV accumulation is at a higher level after 1^st^ injection compared to 5^th^ injection in female rats.
6. Insignificantly higher accumulation of RDV was observed after 5^th^ injection of 387-R-Med in male and female rats.
7. Higher accumulation of RDV was observed in male and female rats, after 1^st^ injection when compared to the 5^th^ injection of 376-R-SBECD (Gilead).
8. For all doses, NV-CoV-2 was detected in rat plasma producing an initial increase that peaked between 4-8 hours, but decreases to below detection level between 24 to 48 hrs.
9. Last but not the least, in all the cases our encapsulated polymer is nontoxic to the animals, based on their steady level of body weight.

## Discussions

RDV formerly known as GS-5734, is a nucleotide analogue that was originally developed as a treatment against Ebola **[3]**. This drug can also inhibit coronavirus replication by inhibiting RNA polymerases (RdRp4). This compound has shown broad antiviral activity *in vitro* against Middle East respiratory syndrome coronavirus (MERS-CoV), severe acute respiratory syndrome coronavirus 1 (SARS-CoV-1) and SARS-CoV-2 **[4-6]**.

In animal studies, RDV has been found effective in protecting rhesus monkeys from MERS-CoV infection when given prior to infection **[8]**. It also protected African green monkeys from Nipah virus, a cause of fatal encephalitis and rhesus monkeys from the Ebola virus **[9, 10]**.

A randomized, well marked, controlled animal study with 12 rhesus monkeys infected with SARS-CoV-2 reported that an attenuation of respiratory symptoms and reduction in lung damage with RDV administered 12 hours after virus infection **[11]**. However, efficacy of RDV *in vitro* or in animals does not match with the clinical outcomes in humans. We searched for other compounds or drugs that could be used in conjunction with RDV to potentiate its effect against SARS-CoV-2 and minimize the side effects of RDV.

Recently we have reported that our lab-made nano-polymer (NV-CoV-2) can encapsulate RDV and guards it from degradation in the blood stream **[15, 16]**. This nanoviricide® bio-mimetic polymer binds and engulfs a virus particle into the polymeric nanoviricide, acting like a “Venus-fly-trap”. Once engulfed, the virus particle gets destroyed **(Figure 11)**.

**Fig. 11:**
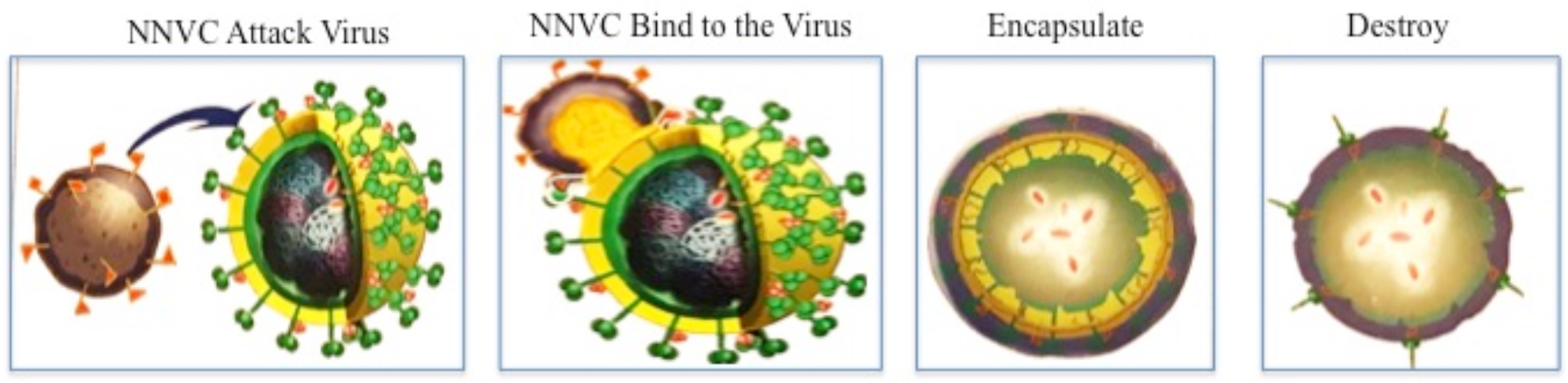
Mechanism of *Nanoviricide* Action

Using a plug-and-play approach, we can change the virus binding ligand portion of this nano-medicine to attack a different virus. We have already tested several drug candidates for broad-spectrum anti-coronavirus effectiveness in cell culture studies. One of the coronavirus strains (h-CoV-NL63) that we studied, uses the same cell surface receptor ACE2 (angiotensin converting enzyme-2) that is shared by SARS-CoV-2 and SARS-CoV-1. Out of our several test drug candidates, NV-CoV-2-R showed as much as 15-times more effectiveness than *favipiravir* against two different coronaviruses (h-CoV-NL63 and HCoV-229E) **[17, 18]**.

Safety and tolerability of that anti-coronavirus drug candidate was studied here in a rat model and found to be safe and well tolerated. Body weights remained constant and there were no clinical signs of immune or allergic reactions such as itching, biting, twitching, rough coat, etc. Furthermore, there were no observable changes in any organs including large intestine or colon on post mortem in gross histology. This non-GLP safety/tolerability study was conducted under GLP-like conditions by AR BioSystems, Inc., Odessa, Tampa, FL. Further microscopic histology and blood work analyses are in progress **[19]**.

RDV (Veklury, Gilead) is supplied as a lyophilized solid containing 100 mg RDV and 600 mg SBECD (sulfobutylether-β-cyclodextrin), with the requirement of being re-dissolved in WFI water by the Pharmacist and injected into infusion fluid (saline) preferably through a sterile filter [20]. It is likely that re-dissolving fully may not complete and filtration may remove undissolved RDV, thus reducing the dose applied. This could explain the discrepancy between RDV controlled clinical trials and the WHO datasets. Cyclodextrins form colloids at high concentrations and allow insoluble drugs to be held in the colloid. However, the colloid dilutes out into the bloodstream quickly and would lead to falling out of the API if it is not held by a true cage-binding mechanism. Thus, protecting RDV from metabolism and keeping it in an encapsulated form is essential if its full potential is to be realized clinically.

## Conclusion

Detection and Quantification of NV-CoV-2 in rat Plasma samples from NV-1067 Toxicology study was done using a validated LC-MS spectrometry as described in the Materials and Methods section. For all doses, NV-CoV-2 was detected in Rat plasma producing an initial increase that peaked between 4-8 hours. The results show that Plasma concentrations decreased to below detection level between 24 to 48 hrs.

## LIST OF ABBREVIATIONS

NV: Nanoviricides
PK/PD: Pharmacokinetics/ Pharmacodynamics
RDV: Remdesivir
NV-387: Nanoviricides-Polymer 387
NV-387-R: Nanoviricides-Polymer 387-Remdesivir conjugate
SBECD: Commercial encapsulating agent SBECD (GILEAD)
SBECD-R: Commercial Remdesivir conjugated with SBECD
PBS: Phosphate Buffered Saline
DMSO: Dimethyl Sulfoxide
RPL: Rat Plasma
MeOH: Methanol
ADME: Absorption, Distribution, Metabolism and Excretion
SD: Standard Deviation

## Authors’ contribution

All the authors contributed equally to prepare this article, read and approved the final manuscript.

## Acknowledgement

We acknowledge all our colleagues, Secretaries for their help during the preparation of the manuscript by providing all the relevant information.

## Ethical Statement

Not applicable

## References

1. Cucinotta D, Vanelli M. WHO Declares COVID-19 a Pandemic. Acta Biomed. 2020; 91(1):157–160. doi: 10.23750/abm.v91i1.9397.

2. Xie Y, Ogah CA, Jiang X, Li J, Shen J. Nucleoside inhibitors of hepatitis C virus NS5B polymerase: a systematic review. Curr Drug Targets 2016;17:1560–76. Doi: 10.2174/138945011766615120912375126648061.

3. Mulangu S, Dodd LE, Davey RT Jr, Tshiani Mbaya O, Proschan M, Mukadi D, et. al. Randomized, Controlled Trial of Ebola Virus Disease Therapeutics. N Engl. J Med. 2019;381(24):2293–2303. doi: 10.1056/NEJMoa1910993.

4. Gordon CJ, Tchesnokov EP, Feng JY, Porter DP, Götte M. The antiviral compound remdesivir potently inhibits RNA-dependent RNA polymerase from Middle East respiratory syndrome coronavirus. J Biol Chem. 2020; 295(15):4773–4779. doi: 10.1074/jbc.AC120.013056.

5. Sheahan TP, Sims AC, Graham RL, Menachery VD, Gralinski LE, Case JB, et. al. Broad-spectrum antiviral GS-5734 inhibits both epidemic and zoonotic coronaviruses. Sci Transl Med. 2017; 9(396):eaal3653. doi: 10.1126/scitranslmed.aal3653.

6. Wang M, Cao R, Zhang L, Yang X, Liu J, Xu M, et. al. Remdesivir and chloroquine effectively inhibit the recently emerged novel coronavirus (2019-nCoV) in vitro. Cell Res. 2020; 30(3):269–271. doi: 10.1038/s41422-020-0282-0.

7. COVID-19 Treatment Guidelines. 2020. https://www.covid19treatmentguidelines.nih.gov/antiviral-therapy/remdesivir/.

8. de Wit E, Feldmann F, Cronin J, Jordan R, Okumura A, Thomas T, et. al. Prophylactic and therapeutic remdesivir (GS-5734) treatment in the rhesus macaque model of MERS-CoV infection. Proc Natl Acad Sci U S A. 2020;117(12):6771–6776. doi: 10.1073/pnas.1922083117.

9. Lo MK, Feldmann F, Gary JM, Jordan R, Bannister R, Cronin J, et. al. Remdesivir (GS-5734) protects African green monkeys from Nipah virus challenge. Sci Transl Med. 2019;11(494):eaau9242. doi: 10.1126/scitranslmed.aau9242.

10. Warren TK, Jordan R, Lo MK, Ray AS, Mackman RL, Soloveva V, et.al. Therapeutic efficacy of the small molecule GS-5734 against Ebola virus in rhesus monkeys. Nature. 2016;531(7594):381–5. doi: 10.1038/nature17180.

11. Williamson BN, Feldmann F, Schwarz B, Meade-White K, Porter DP, Schulz J, et.al. Clinical benefit of remdesivir in rhesus macaques infected with SARS-CoV-2. Nature. 2020;585(7824):273–276. doi: 10.1038/s41586-020-2423-5.

12. Remdesivir (RDV). 2020. https://www.rxlist.com/consumer_remdesivir_rdv/drugs-condition.htm.

13. Adamsick ML, Gandhi RG, Bidell MR, Elshaboury RH, Bhattacharyya RP, Kim AY, et. al. Remdesivir in Patients with Acute or Chronic Kidney Disease and COVID-19. J Am Soc Nephrol. 2020; 31 (7): 13841386. DOI: 10.1681/ASN.2020050589

14. Saleh J, Peyssonnaux C, Singh KK, Edeas M. Mitochondria and microbiota dysfunction in COVID-19 pathogenesis. Mitochondrion. 2020; 54:1–7. https://doi.org/10.1016/j.mito.2020.06.008.

15. Ashok Chakraborty, Anil Diwan, Vinod Arora, Yogesh Thakur, Preetam Holkar, Vijetha Chinige Nanoviricides Platform Technology based NV-387 polymer Protects Remdesivir from Plasma-Mediated Catabolism in vitro: Importance of its increased lifetime for in vivo action. bioRxiv 2021.10.22.465399; doi.org/10.1101/2021.10.22.465399

15a. https://www.news-medical.net/news/20211028/A-novel-approach-to-improve-the-stabilityof-Remdesivir.aspx

16. https://www.accesswire.com/615993/NanoViricides-Has-Engaged-Calvert-Labs-for-Safety-Pharmacology-Studies-of-Its-Drug-for-the-Treatment-of-COVID-19.

17. https://www.accesswire.com/667514/NanoViricides-Announces-COVID-19-Clinical-Drug-Candidate-NV-CoV-2-was-Effective-Against-SARS-CoV-2-Further-Demonstrating-Its-Broad-Spectrum-Pan-Coronavirus-Activity.

18. https://www.biospace.com/article/releases/nanoviricides-announces-covid-19-clinical-drug-candidate-nv-cov-2-was-effective-against-sars-cov-2-further-demonstrating-its-broad-spectrum-pan-coronavirus-activity/.

19. https://www.marketwatch.com/press-release/nanoviricidesdevelops-highly-effective-broad-spectrum-drug-candidates-againstcoronaviruses-2020-05-12.

20. https://dailymed.nlm.nih.gov/dailymed/lookup.cfm?setid=98b7e6bf-2668-4a61-a874-194eb674b15c&version=7#!.

